# Visually guided analysis of single cell proteomics data with SCP-Viz

**DOI:** 10.64898/2026.11.16.767086

**Authors:** Shelby Ruiz, Cameron Braswell, Benjamin C. Orsburn

## Abstract

Single cell proteomics (SCP) is a specialized, but expanding, field of research today. One of many challenges slowing SCP adoption is the inherent instability of the proteome and the changes that can occur before getting cells to a lab for preparation and analysis. Additionally, all current interpretation tools require experienced bioinformaticians to make the step from data to interpretation. Here, we present a visual toolkit that can accept SCP data from any instrument or common data processing algorithm. To demonstrate the capabilities of this tool we apply it to the analysis of dried digests of single human cancer cells following long room temperature storage. Remarkably, proteomic depth of the dried and stored samples analyzed is higher than the original analysis of these same digests on a similar system directly following isolation. Critically, when comparing control and drug treated cells, both the protein abundance and post-translationally modified peptides appear similar in both fresh and stored digests of these single cells. These findings are highly encouraging and, taken together, may help decrease the barriers for entry and collaboration for the field of SCP globally.

## Main

Single cell sequencing (scSeq) techniques are mature and undoubtedly valuable. But the poor correlation between transcript and protein abundance hasn’t improved with technology.^1^ In fact, in single cells, the correlation between transcript and protein abundance appears to be even worse than bulk cell homogenates.^2^ Ultimately, transcript abundance indicates what the cell intends to do, whereas post-transcriptional, translational, and post-translational regulation produce the proteins and peptides that more directly reflect phenotype and cell activity.^3^

Single cell proteomics (SCP) has developed rapidly over the ten years since the first descriptions of unbiased analysis of a small number of human cells.^4,5^ Today, SCP by liquid chromatography mass spectrometry (LCMS) routinely quantifies hundreds to thousands of proteins in single human cells of typical size^6^ using two broad strategies. In multiplexed approaches, peptides from up to 32 single cells can be labeled and combined into a single LCMS run, enabling as many as 1,000 cells to be analyzed per instrument day.^7,8^ Multiplexing enables throughput similar to other sequencing technologies and has demonstrated benefits in quantitative accuracy^9^, but these data are challenged by stochastic sampling and relatively large batch/plex effects.^10^ Label free approaches have therefore become increasingly popular, with the recognized tradeoff that nearly all commercial LCMS hardware is limited to a maximum of approximately 100 cells per instrument day.^11–13^ Choosing between these approaches centers on tradeoffs in proteomic depth, quantitative accuracy, throughput, and the challenge of mitigating batch and plex effects.

These experimental design choices also presume access to suitable instrumentation, including the advantages afforded by automated sample preparation robots, as well as the expertise to analyze or even explore the resulting data. Despite its growth, SCP remains concentrated in a small number of equipped labs, making it challenging to connect researchers with resources. For example, in the context of instrumentation access, recent work has demonstrated that large proteomic changes can occur when living cells are shipped to central facilities, and that the manner of shipment further alters proteomic measurements.^14^ Processing cells into sealed, dried peptide digests may offer a simpler transport or storage alternative, provided these digests remain stable over time. On the analytical side, informatic developments have driven the reliability of SCP nearly as much as method and hardware advancements.^10,15^ However, these tools remain primarily command line driven iterative data transformations which require bioinformatic capabilities beyond that of many proteomics researchers or collaborators. As such, we first developed the Shiny web interface SCP-Viz as a way to simply visualize SCP data from any instrument or data processing software.^16^ Here we present SCP-Viz 2.0, which integrates multiple tools and concepts developed by the single cell and proteomics communities into a flexible, web browser accessible app.

## RESULTS

### Analysis of stored peptide digests using SCP-Viz

To address both SCP accessibility barriers, we evaluated whether the findings of an SCP study could be reproduced using leftover, sealed tryptic digests from the same initial preparation after 17 months of room temperature storage (**Fig. 1a**). Fresh digests of H358 cells treated with the HDAC inhibitor Mocetinostat or DMSO were analyzed on a TimsTOF SCP shortly after cell sorting using tandem mass tag (TMT) data dependent acquisition (DDA).^17^ The remaining unlabeled digests were analyzed after months of room temperature storage using a label free data independent acquisition (DIA) method on a TimsTOF Ultra 2 system. Drawing biological conclusions across labeled and label free datasets remains a pinnacle challenge in the field. Therefore, we chose to not only evaluate the effects of digest storage time but also to demonstrate the power of SCP-Viz 2.0 through sequential analysis of cells from a single preparation whose proteomes were measured by vastly different approaches.

**Figure 1.**
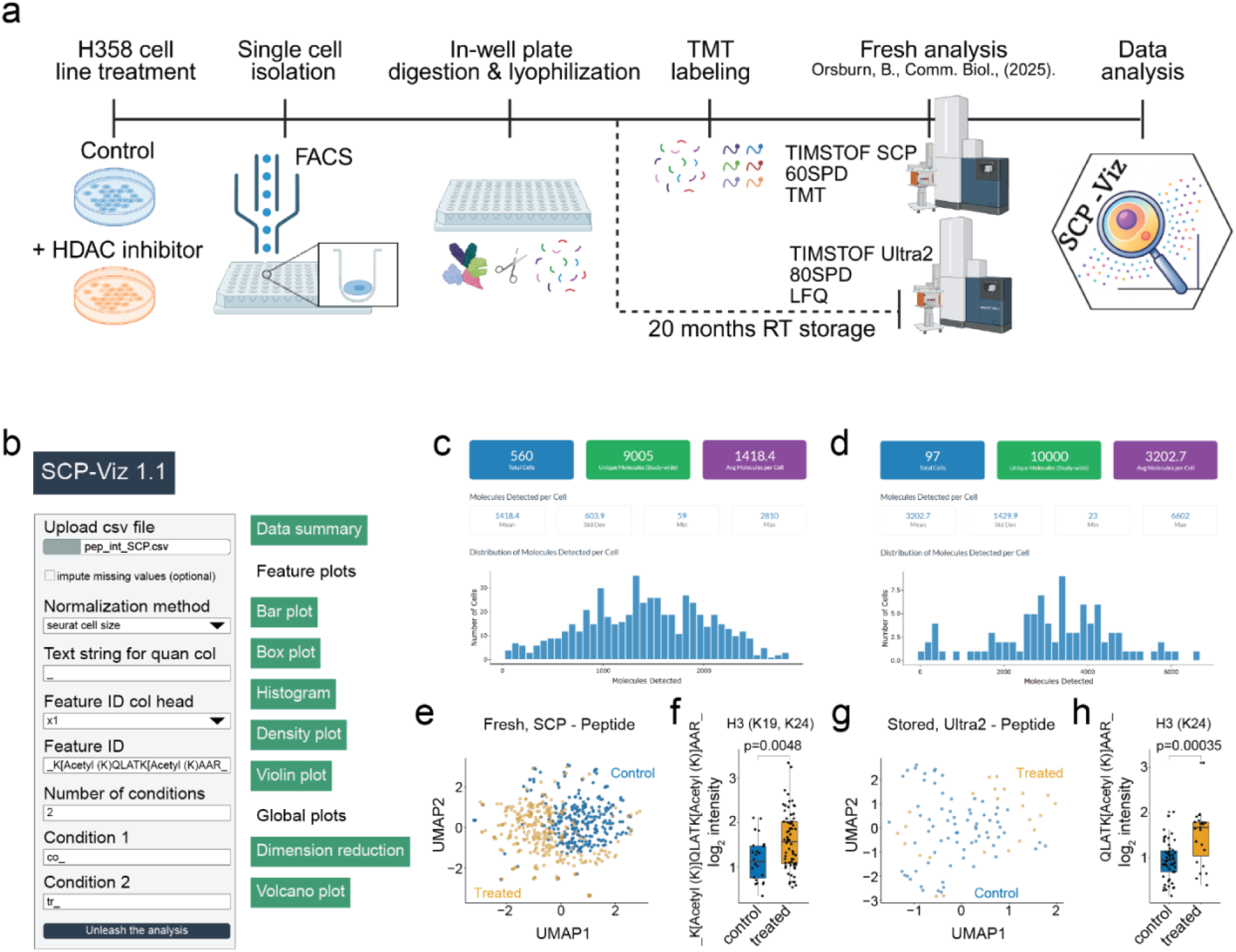
SCP-Viz 1.1 enables comparison of the same cells analyzed with markedly different instrument methods. (**a**) Graphical summary of single cell preparation flow and experimental conditions. (**b**) Within the initial SCP-Viz 1.1 tab, raw or normalized data from a simple database export sheet can be summarized and visualized with the per cell distribution of modified peptide identifications for both (**c**) fresh and (**d**) stored cells. (**e, g**) Dimension reduction plots can easily visualize treatment separation by the selected conditions during upload along with differential abundance testing of selected features of interest for both the fresh (**f**) and stored cell conditions (**h**). Both peptide and protein feature levels can be explored and compared using basic data transformations, plots, and statistics within the SCP-Viz package.

Storage time varies alongside labeling strategy and instrumentation methods, which precludes a direct comparison of storage effects alone. We therefore focused on the retention of the biological ramifications of cancer cell line growth and drug treatment, including single cell heterogeneity within untreated cells and the reproducibility of drug treatment effects. SCP-Viz 2.0 retains all capabilities of the initial public release (**Figure 1b-e**) and we incorporate the updated features throughout to demonstrate the flexibility and depth of peptide and protein data that can be explored at the click of a button.

### Peptide and protein identifications

To evaluate whether long term storage impacted feature detection, we first compared identification rates and overall detection overlap between fresh TMT labeled (SCP) and stored label free (Ultra2) datasets following cell filtering within the SCP-QC tab (**Fig. 2a**). Shared feature detection amongst single cells is inherently heterogenous and features detected in fewer than 20% of cells were common in both datasets. However, these sparsely detected features made up a larger share of the fresh dataset (modified peptides 6460/8986 (71.9%); proteins 999/1692 (59.0%)) than the stored dataset (modified peptides 4530/9659 (46.9%); proteins 573/1777 (32.2%)) (**Fig. 2b**, **e**). We defined features detected in more than 20% of fresh cells as high confidence features, which would be expected to remain detectable at a similar rate following storage. Of these, roughly half of modified peptides but over 90% of proteins were also detected in more than 20% of stored cells. The stored dataset also contained many confidently identified features absent from the fresh dataset, consistent with the greater per cell depth generally achieved by label free single cell methods^18^.

**Figure 2.**
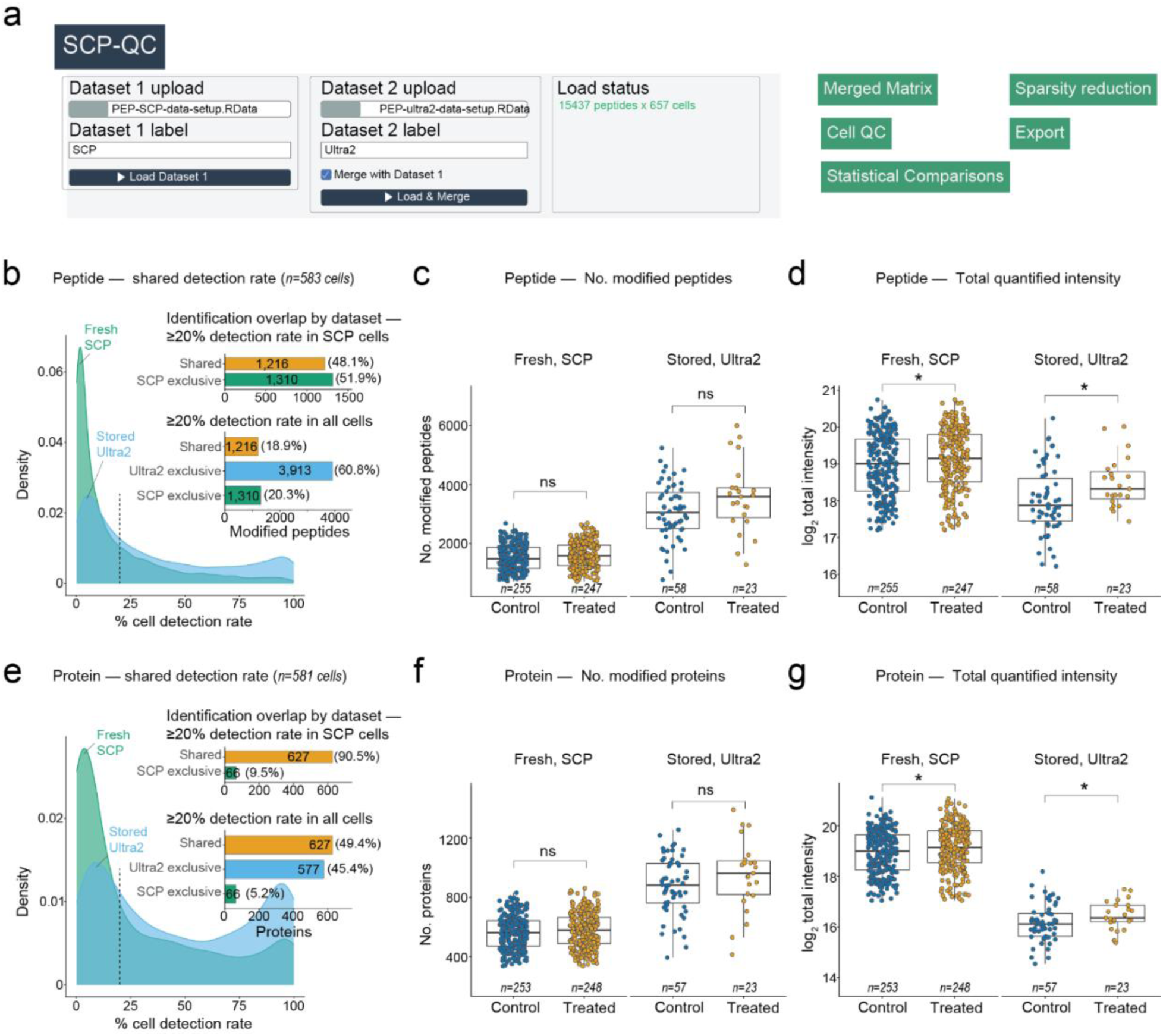
The SCP-QC tab enables direct merging of datasets and the visualization of feature identification rates and total feature quantity comparisons between variables of interest. (**a**) Graphical summary of the SCP-QC tab upload display and visualization subtabs. (**b**) Frequency distribution of the proportion of cells that share detection of specific modified peptides or (**e**) proteins in either dataset. (*inset*) Quantity of features shared or exclusively identified in at least 20% of SCP cells (*top*) or all cells from both datasets (*bottom*) at equivalent detection rate thresholds. (**c**, **d**, **f**, **g**) Box plot distributions of per cell, quantified features and total intensity split by treatment conditions.

Because LCMS feature detection depends on the instrument, acquisition method, search parameters, and each feature’s abundance relative to the detection limit, cross dataset comparisons cannot isolate storage effects. We therefore compared per cell metrics between treatment conditions within each dataset. The number of features detected per cell did not significantly differ between treatment conditions in either dataset (**Fig. 2c**, **f**). In contrast, total quantified intensity at both the peptide and protein level was elevated in Mocetinostat treated cells of the fresh dataset, and this difference was preserved in stored cell digests (**Fig. 2d**, **g**).

### Flexible visual Seurat analysis with SCP-Viz

Beyond visualization, analysis of complex datasets requires identifying, evaluating, and when necessary, correcting or removing technical variables that drive single cell diversity so that true biological heterogeneity can be assessed.^19,20^ The Seurat analysis tab extends the QC tab by quantifying the variance explained by covariates and the incorporation of additional normalization options to the Seurat analysis workflow,^21,22^ including log_2 t_ransformation, median scaling, variance stabilization, and normalization to user-selected reference proteins such as Histone 4 (H4C1) (**Fig. 3a**, **SFig. 1**). Critically, these routes retain missing values as NA rather than replacing them with zeros, enabling visualizations of feature presence versus absence.^23^ The tab also offers several differential abundance tests and post-analysis tools, including p-value distribution histograms, Upset plots comparing significant features across cluster comparisons, correlation of features with any continuous variable, and pathway analysis through Webgestalt integration.^24^ All plots can be exported as PDFs for editing into publication quality figures. Finally, to support reproducibility and collaboration among users of varying experience, each analysis decision is automatically logged to an observable text file and data exploration sessions can be saved and restored to resume analysis at any point.

**Figure 3.**
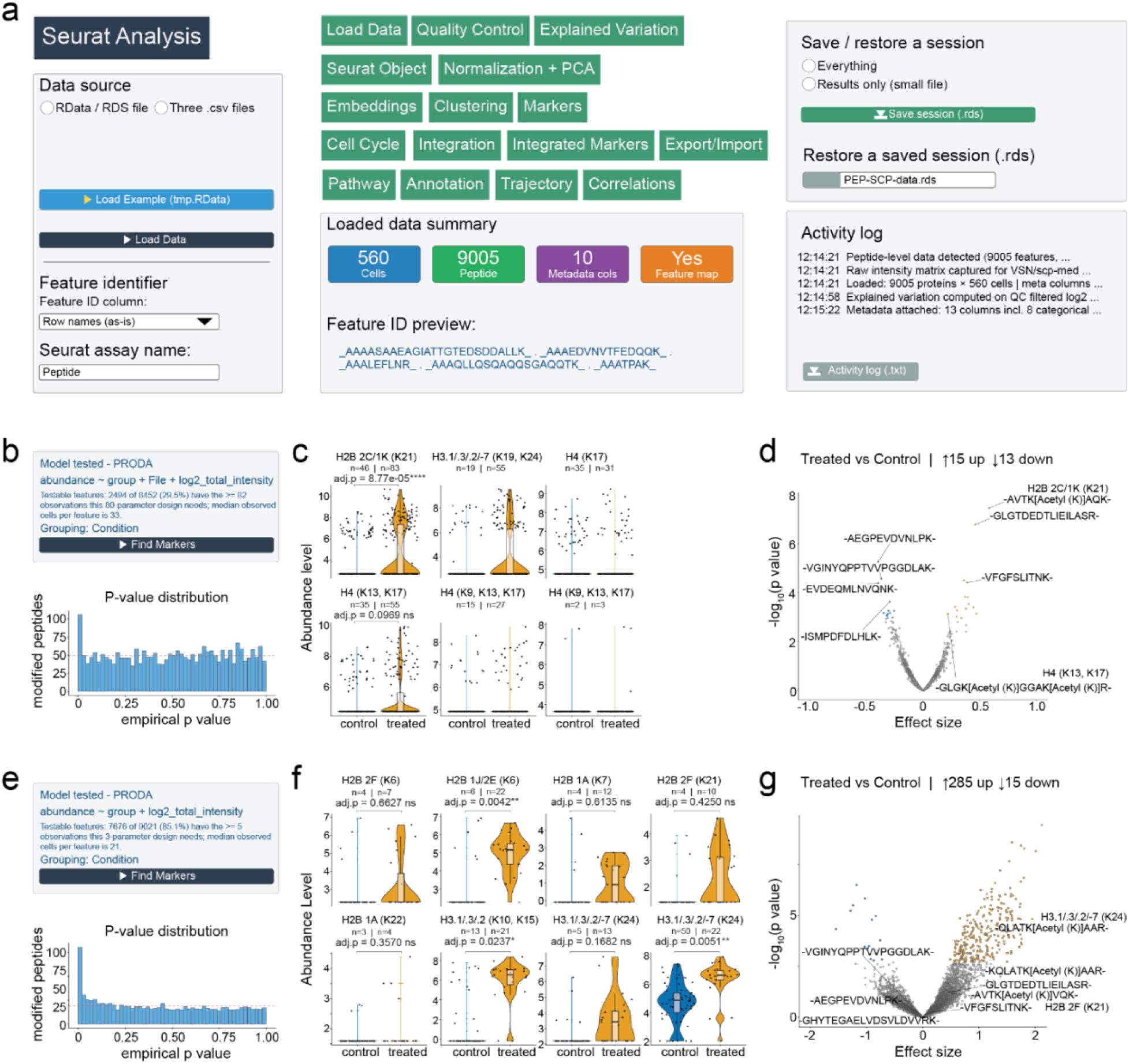
The Seurat analysis tab allows for a full analysis within the Shiny app to identify differentially modified acetylation sites between treatment conditions. (**a**) Graphical representation of the capabilities of the Seurat Analysis tab. (**b** Fresh, **e** Stored) Differential abundance analyses can be performed with a variety of tests including options for covariate modeling inclusion and post analysis checks like p-value distributions. (**c** Fresh, **f** Stored) Scaled abundances by tested treatment conditions can be dynamically selected and visualized as a violin plot, such as peptides of histone acetylation sites. (**d** Fresh, **g** Stored) Volcano plots of the treatment effects can be plotted and modified by the user to include variable labeling selections, color choices, and highlighting of points at specific significance cutoffs (*q<0.1*).

### Analysis of modified peptides

Mocetinostat is a Class I HDAC inhibitor expected to increase histone acetylation, thereby relaxing chromatin structure and inducing transcriptional activation.^25^ As such, we utilized the Seurat analysis tab to evaluate the retention of lysine acetylation site differences between treatment conditions across long term storage.

Histone acetylation sites were largely distinct between datasets (**Table 1**, **Supplementary Table 2**). Six acetylation sites were detected in the fresh digests, compared to eight unique acetylation sites from nine unique peptides in stored digests, for a total of thirteen unique sites. Only a single site at Histone H3 (K24) was identified in both datasets. This limited overlap likely reflects differences in the masses of modified peptides from the lysine rich region of histone tails, as TMT labeling increases the mass of the peptide, enabling smaller peptide ions to be within the scanning m/z window of the instrument.^26^

**Table 1:** Detected acetylation sites across datasets. *This acetylation site was detected in other SCP SPD datasets but not the utilized 60 SPD SCP data in this comparison.

| ***Dataset identified*** | ***Histone acetylation site*** | ***Fresh, TMT SCP peptides*** | ***Stored, LFQ Ultra2 peptides*** |
|---|---|---|---|
| *Ultra2* | H2B 2F (K6) |  | _PDPAK[Acetyl (K)]SAPAPK_ |
| *Ultra2* | H2B 1J/2E (K6) |  | _PEPAK[Acetyl (K)]SAPAPK_ |
| *Ultra2* | H2B 1A (K7) |  | _PEVSSK[Acetyl (K)]GATISK_ |
| *SCP* | H2B 2C/1K (K21) | _[K][TMTpro (N-Term)]AVTK[Acetyl (K)]AQK[TMTpro (K)]_ |  |
| *Ultra2* | H2B 2F (K21) |  | _AVTK[Acetyl (K)]VQK_ |
| *Ultra2* | H2B 1A (K22) |  | _AVVK[Acetyl (K)]TQK_ |
| *Ultra2* | H3.1/H3.3/H3.2 (K10) |  | _K[Acetyl (K)]STGGK[Acetyl (K)]APR_ |
| *SCP\*;Ultra2* | H3.1/H3.3/H3.2 (K15) |  | _K[Acetyl (K)]STGGK[Acetyl (K)]APR_ |
| *SCP* | H3.1/H3.3/H3.2/H3-7 (K19) | _[R][TMTpro (N-Term)]K[Acetyl (K)]QLATK[Acetyl (K)]AAR_ |  |
| ***SCP;Ultra2*** | H3.1/H3.3/H3.2/H3-7 (K24) | _[R][TMTpro (N-Term)]K[Acetyl (K)]QLATK[Acetyl (K)]AAR_ | _KQLATK[Acetyl (K)]AAR_ |
|  |  |  | _QLATK[Acetyl (K)]AAR_ |
| *SCP* | H4 (K9) | _[K][TMTpro (N-Term)]GGK[Acetyl (K)]GLGK[Acetyl (K)]GGAK[Acetyl (K)]R_ |  |
|  |  | _[K][TMTpro (N-Term)]GKGGK[Acetyl (K)]GLGK[Acetyl (K)]GGAK[Acetyl (K)]R_ |  |
| *SCP* | H4 (K13) | _[K][TMTpro (N-Term)]GGK[Acetyl (K)]GLGK[Acetyl (K)]GGAK[Acetyl (K)]R_ |  |
|  |  | _[K][TMTpro (N-Term)]GKGGK[Acetyl (K)]GLGK[Acetyl (K)]GGAK[Acetyl (K)]R_ |  |
|  |  | _[K][TMTpro (N-Term)]GLGK[Acetyl (K)]GGAK[Acetyl (K)]R_ |  |
| *SCP* | H4 (K17) | _[K][TMTpro (N-Term)]GGK[Acetyl (K)]GLGK[Acetyl (K)]GGAK[Acetyl (K)]R_ |  |
|  |  | _[K][TMTpro (N-Term)]GKGGK[Acetyl (K)]GLGK[Acetyl (K)]GGAK[Acetyl (K)]R_ |  |
|  |  | _[K][TMTpro (N-Term)]GLGK[Acetyl (K)]GGAK[Acetyl (K)]R_ |  |
|  |  | _[K][TMTpro (N-Term)]GLGK[Acetyl (K)]GGAK[TMTpro (K)]_ |  |

As expected, Histone H3 (K24) acetylation was detected more frequently and at higher abundances in mocetinostat treated cells in both datasets (**Fig. 3c**, **f**).^26,27^ The fresh dataset contained too few observations of this site for the selected differential abundance model, but a Wilcoxon rank sum test on total intensity normalized values showed a significant difference between treatment conditions in both fresh and stored cells (**Fig. 1f**, **h**). Most other acetylation sites showed higher abundance and detection rates in treated cells without reaching significance (**Supplementary Table 2**). To place these sites in the context of the broader treatment response, we selected them and other significantly differential peptides from either dataset in the Markers tab drop down menu and visualized treatment effects with volcano plots (**Fig. 3d**, **g**).

### Unsupervised clustering of fresh multiplexed single cells

Beyond the large number of repeated measurements and cell level presence versus absence detection, SCP offers a key advantage over bulk proteomics when comparing treatment effects. Drug treated cells in culture may form distinct populations based on their response stage (early, complete, or delayed), drug uptake to size ratios, cell cycle phase at treatment onset, or a simple diversity of cells within a single culture.^28,29^ To identify populations of cells with similar proteomes beyond treatment, we first performed Seurat based clustering analysis on fresh cells.

Clustering requires several parameter choices, including the number of variable features, principal components, and nearest neighbors used for dimension reduction, and SCP-Viz automatically generates diagnostic plots to guide these decisions. Here, we normalized protein level data with SCTransform^30^ and performed principal component analysis on the identified highly variable features (**Fig. 4a**). The explained variance plateaued after approximately ten principal components, and cells distinctly separated by treatment condition along the first two components alone, with the top contributing proteins annotated by direction (**Fig. 4b**-**d**). Clustree analysis visualized how cells partitioned across increasing clustering resolutions and how stable these assignments were, while silhouette scores summarized within-cluster cohesion of cells (**Fig. 4e**, **f**). Rather than separating into treated and untreated groups, cells formed three populations: cluster 0 (C0) and 1 (C1) were dominated by treated and untreated cells, respectively, while cluster 2 (C2) consisted of a relatively even mixture of both conditions (**Fig. 4g**, **h**).

**Figure 4.**
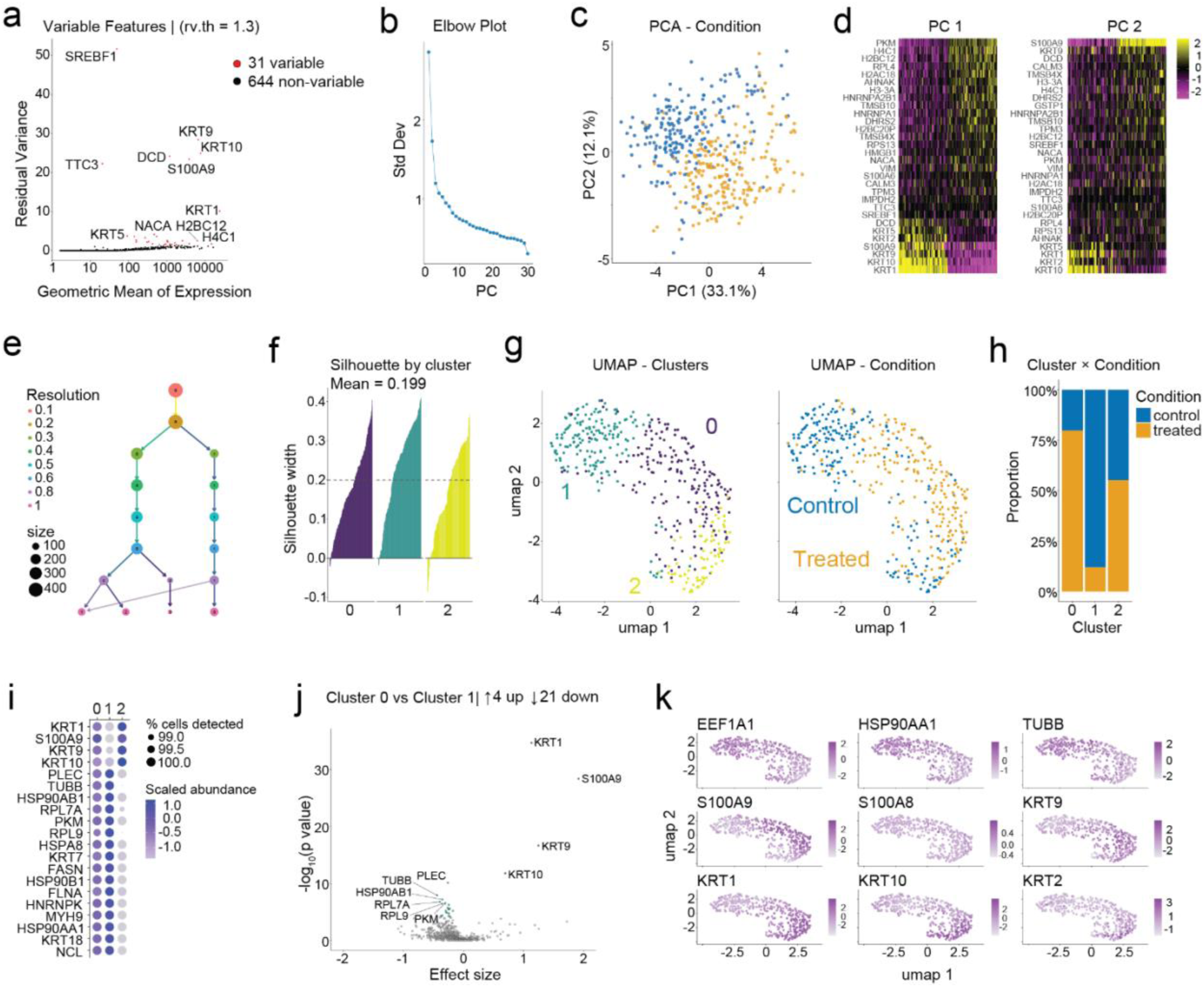
Clustering protein level analysis of Fresh, TMT labeled cells analyzed on a TIMSTOF SCP. (**a**) Highly variable features identified after SCTransform normalization. (**b**) Variance explained per principal component, tapering after PC10. (**c**) Cells separate by treatment condition along PC1 and PC2. (**d**) Top protein drivers of PC1 and PC2, annotated by separation direction. (**e**) Clustree plot showing cluster partitioning and membership stability across resolutions. (**f**) Silhouette scores showing within-cluster cohesion. (**g**) Cells colored by cluster. (**h**) Treatment composition of each cluster; clusters 0 and 1 are biased toward treated or control cells, and cluster 2 is evenly mixed. (**i**) Comparison of protein abundances between clusters 0 and 1, with scaled abundances of the top differential proteins shown across all clusters as a dot plot. (**j**) Volcano plot highlighting selected proteins among all tested proteins. (**k**) UMAP overlay of selected protein abundances across all cells.

To compare this reanalysis with the originally published results^31^, we tested for differential abundance between C0 and C1 (**Fig. 4i**). Scaled abundances of the top differential proteins can be viewed across all clusters as a dot plot (**Fig. 4i**). Additionally, individual proteins can be examined within the context of all tested proteins on a volcano plot or across all analyzed cells as a UMAP feature overlay (**Fig. 4j**, **k**).

Relative to the control dominated cluster (C1), the treated dominated cluster (C0) showed lower abundance of several 60S ribosomal subunit proteins (RPL7A, RPL9, RPL10A), molecular chaperones (HSP90AB1, HSP90B1, HSP90AA1), and proteins involved in cytoskeletal reorganization (PLEC, FLNA, MYH9). The treatment dominated cell cluster showed significantly higher abundance of select keratins (KRT1, KRT10, KRT 9) and the multifunctional, regulatory protein S100A9. Lower ribosomal protein abundance is consistent with reduced translational capacity, and KRT1 and KRT10 are tightly coregulated markers of cells in the early stages of differentiation^32^. These proteomic alterations suggest that mocetinostat treatment may shift cells toward a more differentiated state. Comparisons between C0 and the mixed treatment cluster (C2) identified similar differentially abundant proteins but with increased effect sizes relative to untreated dominated cluster (C1) (**Supplemental Table 4**).

### Exploration of coregulatory relationships between clusters and treatment conditions

Keratin expression can be lineage specific and several keratins, including KRT7, KRT8, KRT18, and KRT19, serve as markers of lung adenocarcinoma.^33^ Keratins are also common proteomic contaminants, particularly the epidermal keratins expressed on human skin KRT1 and KRT10. However, keratin expression also shifts when epithelial cells such as H358 transdifferentiate.^32^ This keratin switching occurs during adenocarcinoma-to-squamous cell carcinoma transformation (AST), a documented phenomenon of treatment resistance in cancers carrying KRAS G12C mutations,^34,35^ which H358 cells harbor.^36^ Cells undergoing AST lose adenocarcinoma markers such as KRT7 and KRT18 and gain squamous markers such as KRT5 and KRT6A.^34^

To distinguish keratin switching from contamination, we compared the abundance of top differential proteins and selected keratins across clusters and treatment conditions (**Fig. 5a**, **b**). KRT5 and KRT6A abundance increased across the identified clusters, alongside KRT1 and KRT10, although these differences were not significant. Over the same clusters, KRT7 and KRT18 abundance decreased significantly, with a slightly stronger effect in mocetinostat treated cells within each cluster. Because contamination can add keratin signal but cannot reduce endogenous keratins, this coordinated loss of adenocarcinoma keratins supports a biological shift rather than a contamination artifact. These trends were also observed in the stored dataset (**Fig. 5c**, **d**) and the differential abundance results across tests and datasets can be found in **Supplemental Table 4**.

**Figure 5.**
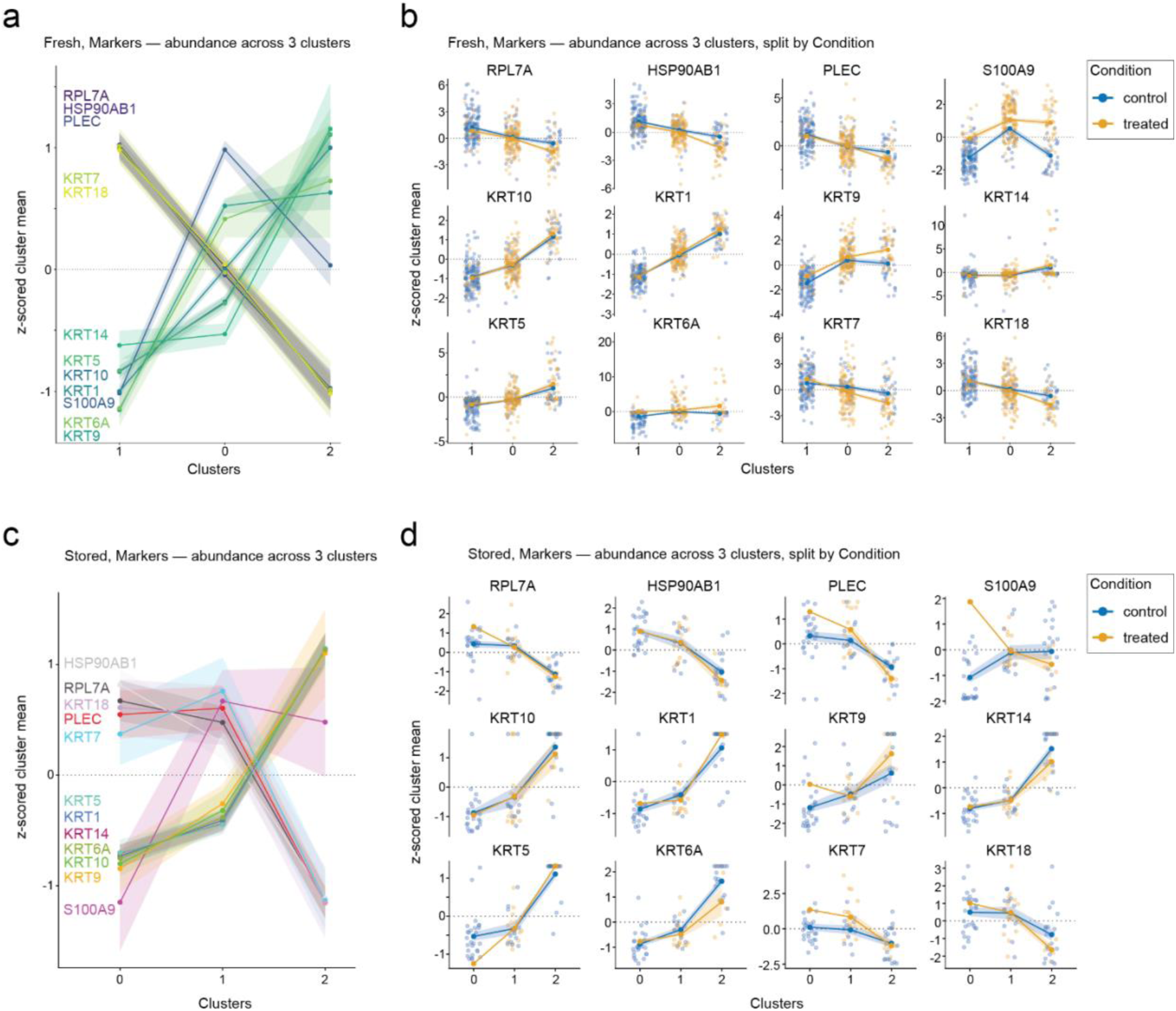
Select protein abundance analysis across cluster and within treatment conditions. Abundance trends of top differential proteins and selected keratins across clusters (**a**) and treatment conditions (**b**). Squamous-like keratins (KRT1, KRT5, KRT6A, KRT10) increase across clusters, coinciding with a significant decrease in KRT7 and KRT18, with a slightly stronger effect in Mocetinostat-treated cells across clusters. (**c**). Individually plotted protein abundances split by condition (**d**).

Although SCP-Viz does not yet include a formal statistical test for these relationships, visualizing coregulated abundance patterns across clusters and metadata for individual cells is a powerful opportunity to explore the strength of specific hypotheses. Together with the overlapping differential proteins between cluster comparisons and the mixed treatment composition of C2 cells, these patterns suggest that mocetinostat promotes a state of epithelial cell differentiation that a subset of untreated cells may acquire spontaneously.

### Clustering analysis of S, label free analyzed single-cell digests

While the acquisition methods of fresh and stored treated and untreated cells presents an imperfect comparison, we can leverage the increase in relative protein identifications from LFQ methods to examine a relatively small number of stored cells. As expected, clustering of the stored cells yielded distinct but sparse, fractured clusters (**Fig. 6a**, **b**).^37^ However, as in the fresh dataset, we observed three cell populations with similar treatment dominated or mixed cell populations with relatively distinct upregulation of specific markers (**Fig. 6c**). Compared to the first untreated dominated cluster (C0), the treatment dominated cluster (C1) demonstrated an upregulation in the levels of the proinflammatory cytokine IL18, the stress associated proteases CTSC, CAPN2, and extracellular matrix, pro-migratory associated proteins HAS1, LGALS3, LIMA1, and ACTBL2 (**Fig. 6d**). Upregulation of these stress responsive proteins was accompanied by increased keratinization and downregulation of cell cycling proteins, including those that regulate DNA replication (PCNA, MCM2, MCM3) (**Fig. 6e**). This decreased proliferation signal and increase in promigratory proteins was also observed when the treatment dominated cluster was compared with the other untreated cluster that had slightly more treated cells than the first untreated population (**Fig. 6f**, **g**). Unique from the other untreated dominated cluster, Cluster 2 was significantly enriched in several proteins characteristic of squamous epithelia, including KRT5, KRT6A, NCCRP1, and KPRP along with genes nearby or within the chromosomal region of the epidermal differentiation complex such as S100A9 and ECM1 (**Fig. 6c**).^38^ This recapitulates lineage plasticity findings as in the fresh data and highlights that these untreated cells may exist in culture as a gradient of tumor populations with some differentiated towards a squamous lineage.

**Figure 6.**
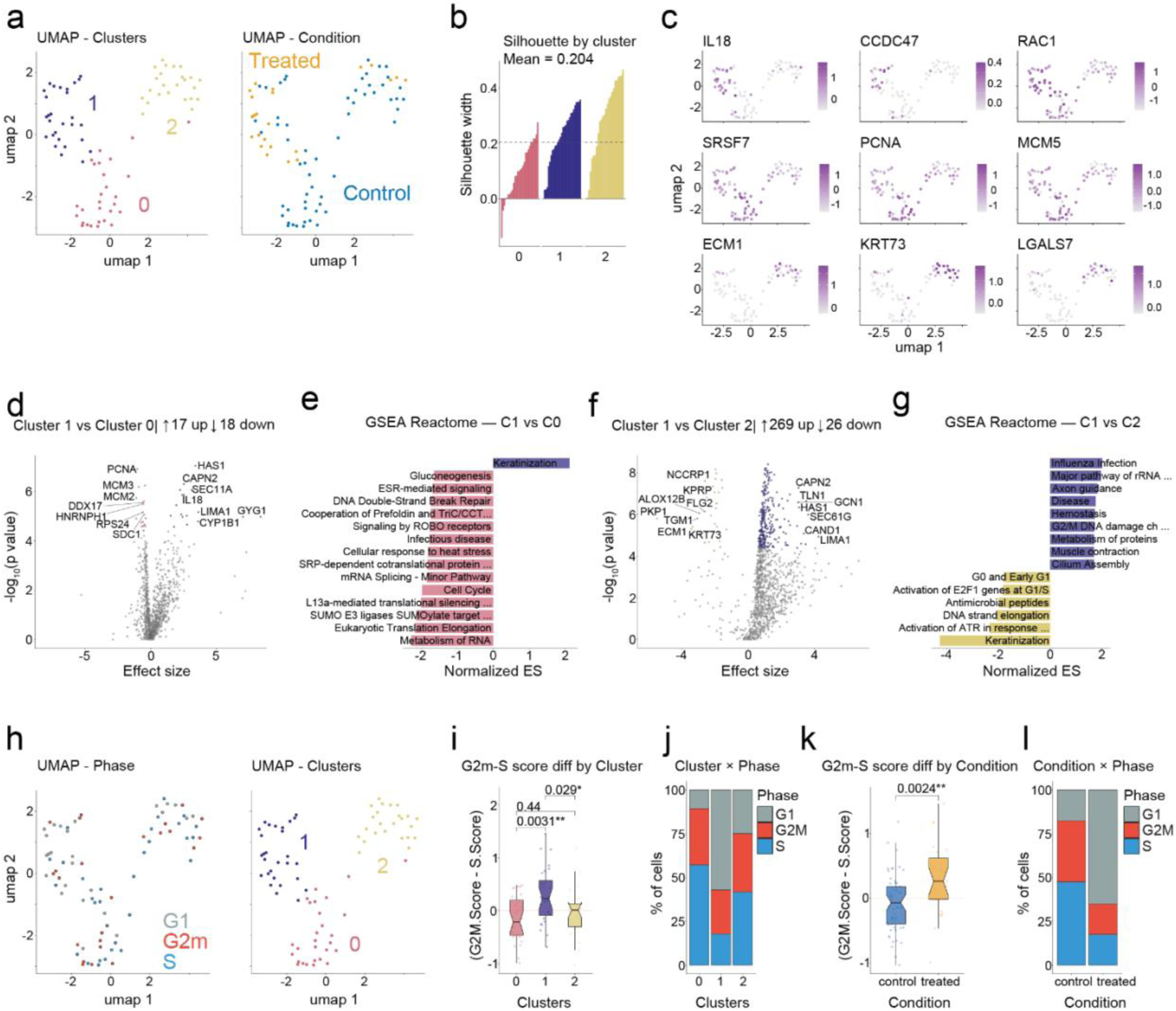
Clustering protein level analysis of stored, LFQ cells analyzed on a TIMSTOF Ultra2. (**a**, **b**) Clustering of stored cells yields distinct but sparse clusters (**a**), shown by treatment condition (**b**). (**c**) Three populations with treatment-dominated or mixed composition and distinct marker upregulation, including squamous markers in cluster 2. (**d**, **e**) Differential protein abundance (d) and pathway enrichment (**e**) for the treatment-dominated cluster (C1) versus the untreated-dominated cluster C0, showing increased stress, protease, and pro-migratory proteins and keratinization, with decreased cell cycle proteins (PCNA, MCM2, MCM3). (**f**, **g**) Differential abundance (**f**) and enrichment (**g**) for C1 versus C2. (**h**-**j**) Cell cycle phase composition (**h**), phase scores (**i**), and G2M minus S score difference (**j**) across clusters. (**k**, **l**) Cell cycle phase composition (**k**) and phase scores (**l**) by treatment condition.

HDAC inhibitors classically induce G1 arrest.^25,27,39,40^ To examine if clusters and treatment conditions could recapitulate this expectation, we utilized the cell cycle annotation features of the Seurat package to bin cells into discrete phases. The treatment dominated cluster was primarily composed of cells exhibiting a G1 phase signature and the difference between G2M and S phase scores was significantly greater than either of the untreated dominant populations (**Fig. 6h-j**). A direct comparison between treated and untreated cells also provides evidence of drug-induced G1 arrest (**Fig. 6k**, **l**). However, proportional differences in cell cycle phases or phase scores between clusters or treatment conditions were not observed in the fresh dataset (**Supplementary Figure 3**). A significant increase in the proportion of cells with a G1 signature only in the stored dataset is unlikely to be due to peptide digest storage. This is more likely to be related to use of cell size normalized but not plex-effect corrected abundances during phase classification, or due to a reduction in the number of identified cell cycle marker proteins (S Phase: 7/43; G2M Phase: 5/54) compared to the stored, LFQ dataset (S Phase: 10/43; G2M Phase: 9/54).

## DISCUSSION

SCP is a field of research that requires specific instrumentation and analytical skills to completely execute an experiment. As such, many SCP experiments may require relatively large collaborative efforts. Work in our group generally requires three groups – the collaborators with the cells, the collaborators or core that can live sort the cells, and our group which ultimately prepares and analyzes the cells. Here, we critically present evidence that basic speedvac drying of peptide digests is an efficient and stable long-term storage method for single cell analysis. Given the fact that peptide digest standards are shipped dry around the world to proteomics labs, the results of this current study may not be altogether surprising. However, the results presented in this technical note seem worth sharing given the literature that has sought to stabilize live cells for shipment and analysis^14^. The authors fully acknowledge that the supporting evidence of stored cell digest stability is limited by the lack of a direct reanalysis because fresh cells were multiplexed on a TIMSTOF SCP system while stored cells were later analyzed on a TIMSTOF Ultra2 system. Due to the corresponding author’s lab relocation and instrument upgrade, an exact replicate analysis between fresh and stored cells is impossible. Further work using single cell digests stored for analysis on the same instrument would be advantageous for separating instrument and methodological related artifacts. Ultimately, treatment effect similarities and the recapitulation of untreated cell population heterogeneity support the idea that single cells can be digested anywhere in the world and delivered to our, or other SCP labs for robust analysis. These findings are highly encouraging for extending the geographic and chronological range of studies in this emerging field.

Additionally, on the point of extending accessibility within SCP, we establish the utility of the updated SCP-Viz 2.0 package, a robust, flexible tool for single cell visualization and analysis. Critically, best data practices follow similar principles^15,41^ but the specifics of each analytical approach should vary in accordance with the implemented instrumentation methods (TMT vs LFQ) or the biological question of interest. For example, TMT based single cell proteomics analyses contain lower levels of missingness between samples of the same plex but induce a large plex effect.^10,41^ If formal normalization is to be avoided, as suggested by others^15^, a high number of observations must be met to allow for the testing of differential features because of the number of plexes.

Furthermore, while some labs may be interested in exploring proteomic differences in relation to cell size, cell size normalization is often paramount to meaningful cell-to-cell comparisons. Cell diameter measurements obtained from capable sorters during dispensing^42^ or calculated size factors inferred from total, mean, or median feature abundances are useful normalization metrics. But as instrument availability varies between labs and methods often differ in the ion population measured, total quantifiable measurements may not accurately reflect cell size. Cell size has been shown to be significantly correlated with the abundance of a specific histone subunit, H4C1, and this quantitative feature may serve as a suitable correction factor for a multitude of single cell studies.^43^ However, H4C1 is not always a useful correction factor. One example is during the single cell analysis of hepatocytes, which are uniquely heterogenous in their ploidy states and whose histone abundance does not linearly scale with polyploidization.^44,45^ This precludes H4C1 abundance as a reliable measurement capable of disentangling protein abundance from the additional various technical factors associated with low input sample preparation loss. Therefore, a versatile range of analysis and normalization options appear necessary to address diverse experimental SCP designs.

Single cell proteomic datasets are full of rich, heterogenous information that often becomes distilled into large cell clusters and a few, static figures for publication. This static *one-and-done* format reduces the availability of the plethora of molecular information available to the field and restricts second party evaluation or primary reanalysis using updated best practices in the field. For example, DIA based proteomics is the main instrumentation method for acquiring single cell proteomic data. There are continuous advancements in database spectral matching,^46,47^ including the incorporation of modified peptide identifications in single cell protein datasets. Furthermore, complex issues still surround quantitative challenges related to peptide rollup and the current approaches to missing values and how to best test for differential abundance when overlapping identifications are sparse and true missingness (presence vs absence) is expected at the protein level for unique cell classes.

Therefore, in contrast to the creation of new analytical methods, we incorporated a multitude of recommended analysis steps (expanded upon by other publications^15,48–50^) and integrated existing packages into a flexible analysis platform built to visually explore and perform robust analyses at the click of a button. This framework supports users of varying levels of analysis expertise and allows for the continued addition of novel, best practices in the field to be incorporated. Furthermore, by providing a range of tools we promote discussions on the analytical methods employed in single cell analysis, welcome feedback on its analytical options, and offer a path towards continued exploration of the multitude of single cell datasets already published.

## ONLINE METHODS

### Treatment of cells

NCI-H-358 cancer cells (H358) were obtained from ATCC and grown in media according to all vendor recommended protocols. Single control cells were treated with DMSO or with 5 µm Mocetinostat (SelecChem S1122) prepared in DMSO. Treated cells were cultured alongside control cells for 24hours prior to rapid washing of cells with ice-cold magnesium and calcium-free PBS (Fisher). The adherent cells were briefly rinsed in 3 mL of 0.05% Trypsin plus EDTA solution (ATCC 30-2001). This solution was rapidly aspirated off and replaced with 3 mL of the same solution. The cells were examined by light field microscopy and incubated at 37 °C with multiple examinations until the adherent cells had lifted off the plate surface. The active trypsin was then quenched by the addition of 7 mL of the original culture media. The 10 mL solution was transferred to sterile 15 mL Falcon tubes (Fisher) and centrifuged at 300 × *g* for 3 minutes to pellet the cells. The supernatant was gently aspirated off, and the cells were resuspended in PBS solution without calcium or magnesium with 0.1% BSA (both, Fisher Scientific) at 1 million cells per mL as estimated by bright field microscopy. Cells for single cell aliquoting were gently dissociated from clumps by slowly pipetting a solution of ∼1 million cells through a Falcon cell strainer (Fisher, 353420) and the cells were placed on wet ice and immediately transported to the JHU Public Health sorting core. Non-viable cells were labeled with a propidium iodide (PI) solution provided by the core facility and briefly vortexed prior to cell isolation and aliquoting. Isolated single cells were deposited directly onto microwell plates containing two microliters of LC-MS grade acetonitrile per well. Cell sorting was performed in early November of 2023 and dried cell lysate plates were stored at 4°C until June of 2024. Plates containing single cells were removed from −80 °C in small batches and were immediately unsealed and placed onto a 95°C heatblock for ∼90 seconds to fully lyse the cells and to dry off the remaining acetonitrile. The plates were cooled to room temperature, and the lysates were digested with a solution of 100 mM TEAB, 0.1% DDM, and 2 ng/µL sequencing grade trypsin. Two µL were added to each blank and single cell well, and 4 µL was added to each carrier channel well. The plates were briefly centrifuged to remove air bubbles and were tightly sealed with plate sealing tape. The digestion was allowed to proceed at room temperature overnight. Following digestion, the plates were thoroughly centrifuged and dried by speedvac.

### Cell storage and LCMS analysis

Dried single cell digests were transported to long term air conditioned storage in June of 2024 where they remained until February of 2025 when they were transported to the University of Pittsburgh and left in an unpacked box. In April of 2025 the first plate of single cell peptide digests were resuspended by placing 20 µL of 0.1% formic acid in water (buffer A) in each well. The digests were briefly vortexed and remained at room temperature for 30 minutes before being loaded into EvoTips following the vendor protocol. The cells were analyzed from 8 separate plates between 4/9/2025 and 4/11/2025. Peptides were separated on an EvoSep One system using a modified 80SPD “WhisperZoom method” in which a 10cm x 75 µm with 1.9 µm particle (PepSep, now Bruker Daltronic) was used in place of the vendor recommended 5cm column. The column was connected to a 10 µm CaptiveSpray emitter by a “zero dead volume” junction. Peptides were analyzed with a TIMSTOF Ultra2 system using the default diaPASEF low sample load method provided by the vendor. For analysis, all Bruker .d files were converted to .HTRMS with the free Biognosys HTRMS converter tool and were analyzed in SpectroNaut 21. All vendor default workflows were used with the exceptions that alkylation of cysteines were not employed, lysine acetylation was added as dynamic modification and a maximum of 4 missed cleavage events were considered.

### Label free quantification data analysis

For database searches, MS .d raw files were converted to .htrms files and searched using Spectronaut v21.0 (Biognosys). Files were processed via the command line in library-free directDIA+ mode against the Reviewed Uniprot human database (20,431 entries, updated on October 26, 2023). Spectronaut parameters are specified in Table 2. Data was exported for down-stream processing at the peptide and protein group level.

**Table 2.**
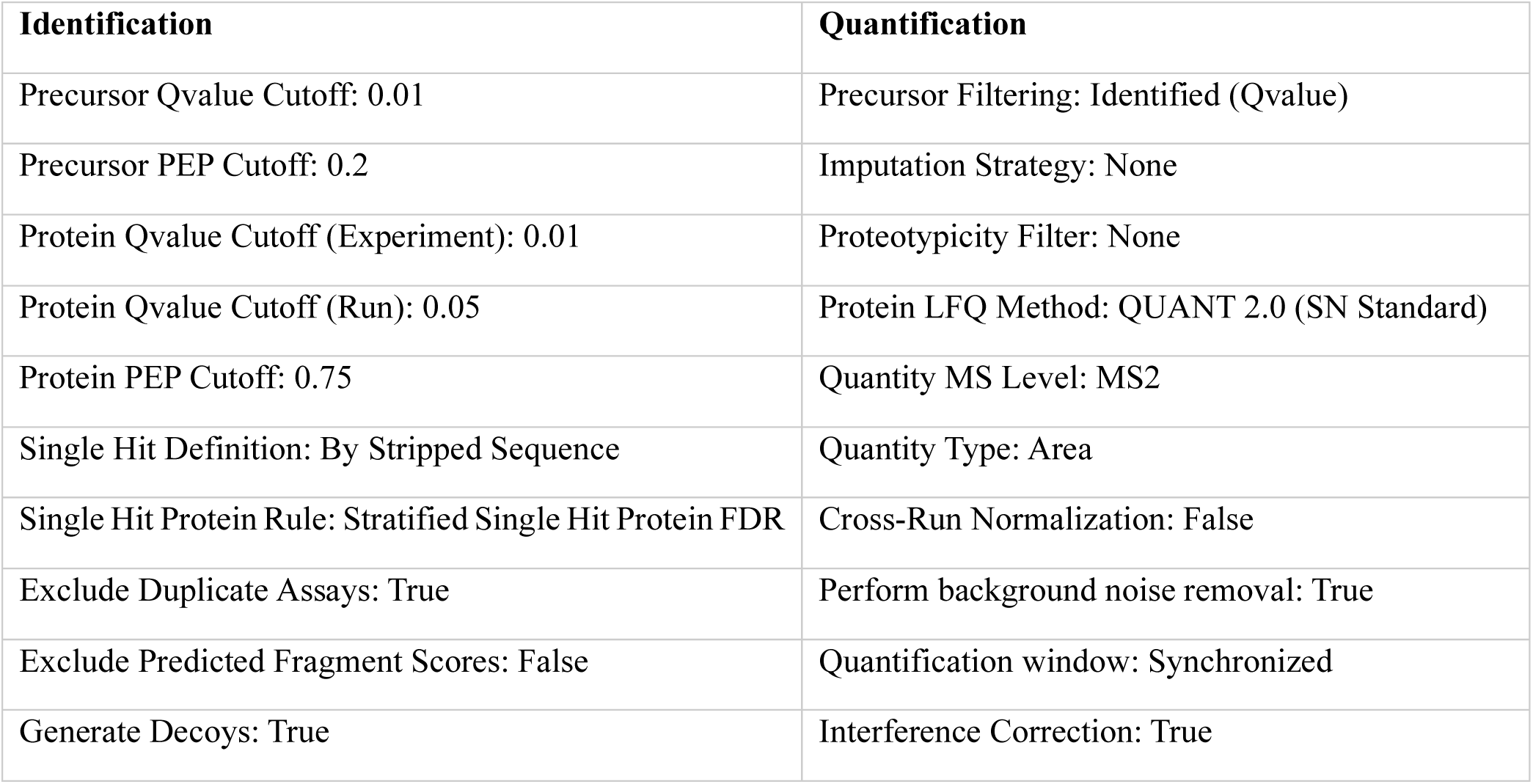
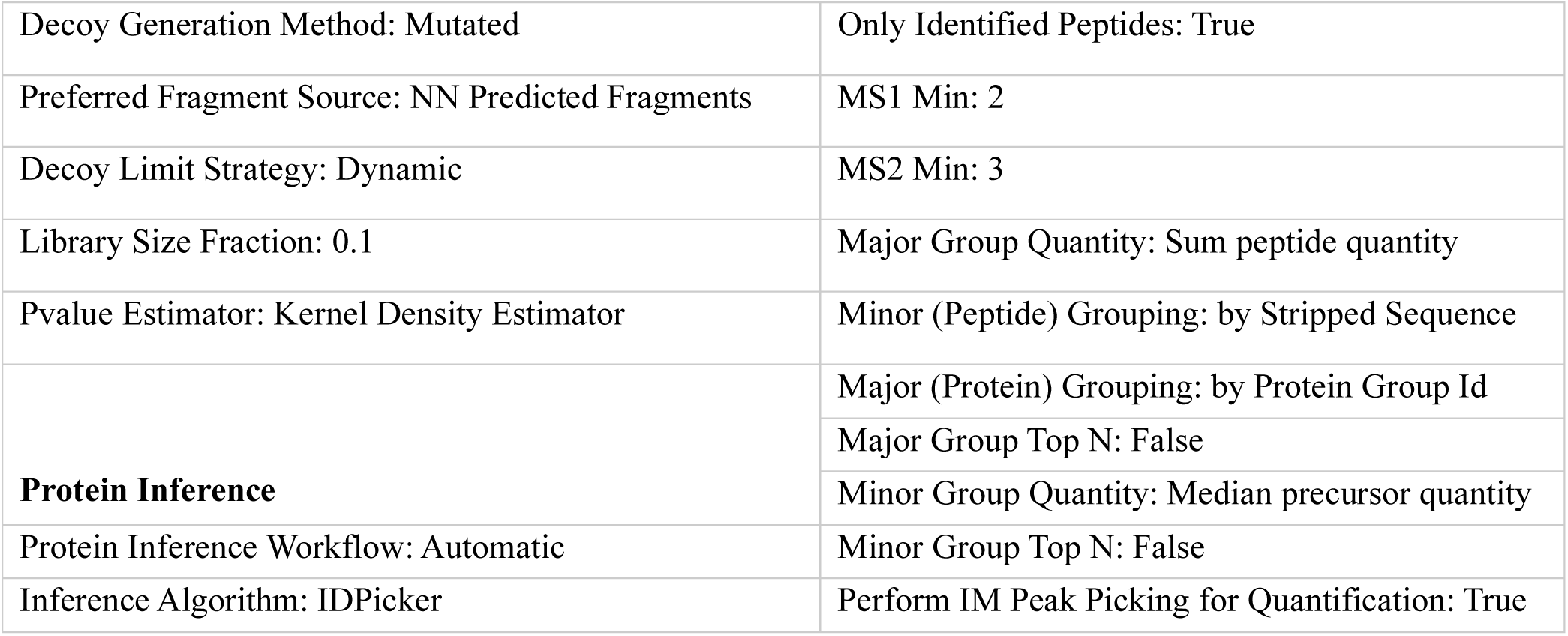
Spectronaut search parameters.

### Data Interpretation

Unnormalized peptide and protein pivot reports were exported from SpectroNaut and Proteome Discoverer and converted in R to .csv file or .RData input for analysis in SCP-Viz. To facilitate R processing, the sample headers can also be cleaned within excel to remove brackets, periods and commas prior to analysis in SCP-Viz on the University of Pittsburgh HPC environment. For comparisons at the modified peptide level, all TMT modifications were removed from the annotated sequence while all other peptide modifications were retained. For the protein level input only unique peptides that mapped to one protein were retained for analysis. All raw data input is formatted for SCP-Viz input and can be found in **Supplementary Table 1**.

Figure 1: Raw modified peptide input was loaded into the SCP-Viz 1.1 tab and visualized following the Seurat cell size normalization method, which is equivalent to log(1p) feature intensity / total cell intensity * 10,000. The pca plot was created with the following selections: 10% features quantified, log2 transformation, and n.neighbors=15.

Figure 2: Raw peptide and protein datasets were loaded into the SCP-Viz SCP-QC tab and merged. Cells were grouped by dataset and passed quality control filtering (n=583/581 cells) if at least 500 modified peptides or 200 proteins were quantified and the cell’s total intensity fell within two standard deviations of the distribution mean. Overlapping features were evaluated at a cutoff of 20% present call across all cells in both datasets. Total feature identifications and total intensities were compared between cell treatment conditions using Dunn’s test with Bonferroni correction (**Supplementary Table 2**).

For treatment effect comparisons, single cell datasets were individually loaded into the Seurat analysis tab and analyzed using the selection parameters referenced in **Supplementary Table 3** and **4**. In brief, cells were filtered as in the SCP-QC tab. For protein level analyses, a 20% sparsity reduction of quantified proteins was employed. For the modified peptide feature level datasets, filtered cells were log_2 n_ormalized and corrected for log_2(_total intensity) and File/Plex if applicable. The top 1000 variable features were used for PCA with nipals, a non-linear iterative partial least square model that allows for missing values. TMT Channel overlay of the Fresh, SCP TMT dataset identified unusual clustering of the 133C Quan channel. Therefore, all 133C labeled cells were removed from both feature level analyses. Differential peptide abundance testing between treatment conditions was performed with ProDA^51^ and log_2 t_otal intensity and File were included in the model as covariates. Filtered protein level data was normalized with SCTransform v2^52^ and corrected for File in the TMT data. Differential abundance testing between clusters or treatment conditions was performed with Seurat’s FindMarkers() or FindAllMarkers() function using the Wilcoxon test. GSEA pathway analysis was performed with WebGestalt R functions integrated into the app.^24^ All figures were exported directly from SCP-Viz as pdfs and briefly edited in adobe illustrator with minor modifications to text size and visual arrangements.

## Data Availability

All .raw and processed data are publicly available on the ProteomeXchange partner repository, www.Massive.UCSD.edu.^53^ TIMSTOF SCP files and library files are available as accession PXD051994 and 17-month room temperature stored digests are available as accession PXD079517. All data presented in this manuscript has been loaded as example data which can be automatically loaded into the SCP-Viz 2.0 local or web interfaces provided below.

## Code Availability

The SCP-Viz 2.0 Shiny app is web accessible at https://shiny.crc.pitt.edu/. The app can also be run locally and downloaded along with accompanying example data and user guides at https://github.com/OrsburnLab-Pitt/SCP-Viz-2p0.

## Artificial intelligence use statement

Claude was used to turn user written scripts into a functional shiny app and iterated on with the user over multiple sessions. All the code has been verified by the user to produce the plots as intended. Further details can be found in the user guide for detailed pipelines and functions.

## ASSOCIATED CONTENT

**Supporting Information**. The following files are available free of charge.

Supplemental Table 1. All feature files formatted for SCP-Viz input.

Supplemental Table 2. Histone acetylated peptide mapping to sites and Dunn’s test results from the SCP QC tab.

Supplemental Table 3. Modified peptide analysis parameters and differential abundance results for both the fresh and stored datasets.

Supplemental Table 4. Protein analysis parameters, differential abundance results, pathway analysis results for both the fresh and stored datasets.

## ACKNOWLEDGMENT

We would like to thank Dr. Ronnie Cutler and Dr. Simone Sidoli for sharing Skyline templates for the targeted analysis of histone PTMs. We would also like to thank Ahmed Warshanna, for his extensive notes and comment lines included in the original SCP-Viz 1.0 build.

