## Supplementary material for "Visually guided analysis of single cell proteomics data with SCP-Viz": S. Fig 1 and 2

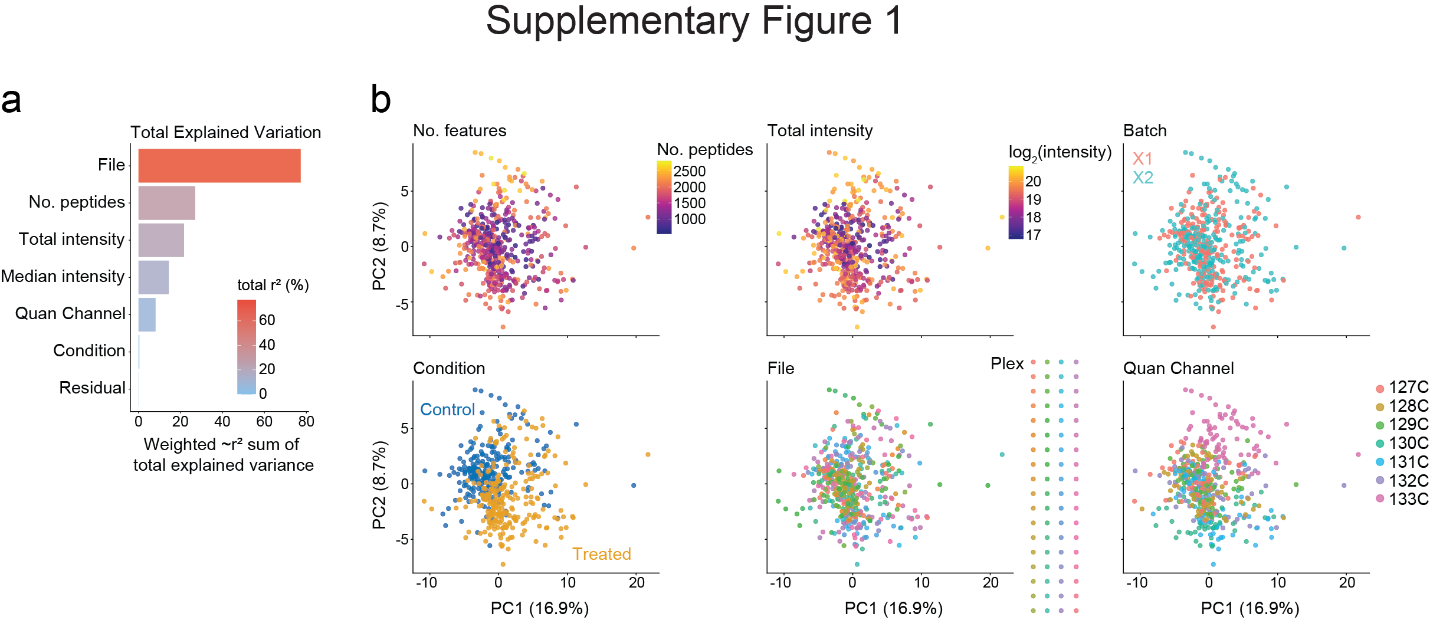

**Supplementary Figure 1.** Identification of prominent and confounding technical variables. (**a**) Selectable covariates of interest were tested for explained variation revealing File/Plex effect as the strongest driver of single cell heterogeneity. (**b**) Following log_2_ normalization and covariate correction of log_2_ total intensity and File, the cells were visualized and overlaid with other technical variables following principal component analysis. Here, the 133C TMT Quan Channel clustered by itself and did not separate by treatment condition. This represents a confounding variable and all cells labeled with the 133C channel were removed from both feature level analyses.

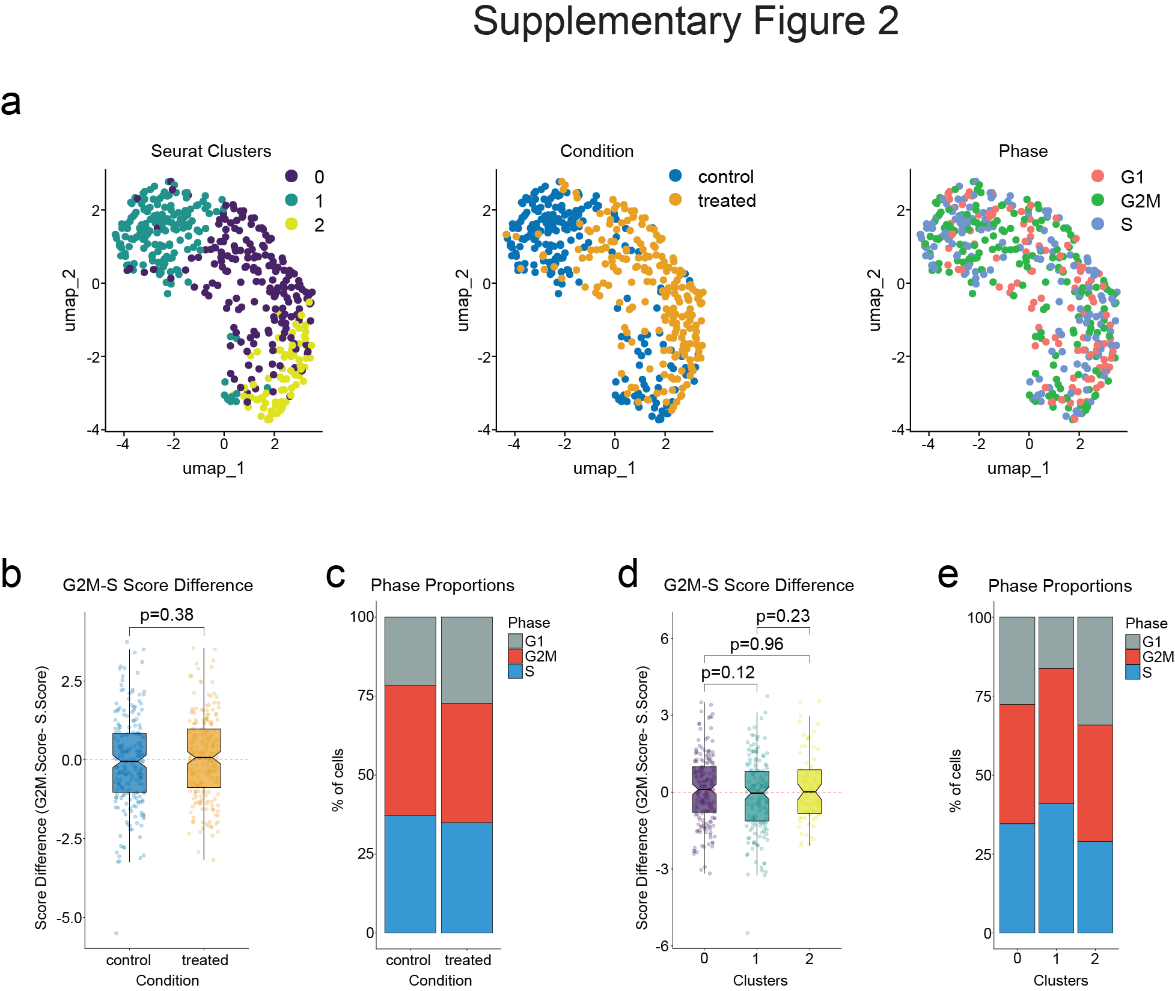

**Supplementary Figure 2.** Cell cycle phase classification differences across Fresh, TMT labeled cells. (**a**) Dimension reduction demonstrated that the identified clusters of fresh cells broadly separated by condition but not a particular cell cycle phase. Despite significant differences in cell cycle phase composition of treatment conditions and clusters identified in the stored dataset (**b**) G2M minus S score differences and (**c**) phase proportions did not significantly differ by condition or (**d, e**) across clusters in the fresh dataset.
